# Sequential Use of Two Capsule-Targeting Klebsiella Phages Reveals Order-Dependent Efficacy and Distinct Resistance Pathways *in vitro* and *in vivo*

**DOI:** 10.64898/2026.06.10.731307

**Authors:** Zhanybek Selpiev, Patrycja Olszewska, Bartłomiej Grygorcewicz, Sebastian Leptihn, Belinda Loh

**Author notes:** Corresponding author: Dr. Belinda Loh, Fraunhofer Institute for Cell Therapy & Immunology (IZI), Department of Infection Research & Diagnostics, Antimicrobial Biotechnology Unit, Perlickstr. 1, Leipzig, 04103, Germany.

## Abstract

Bacteriophage cocktails are widely assumed to improve treatment of multidrug-resistant *Klebsiella pneumoniae*, yet many therapeutic phages target capsular polysaccharides (CPS), potentially promoting antagonism and shared resistance. Here, we investigated how receptor usage, resistance evolution, and dosing order influence the activity of two K1-specific phages, Loop and Spear, against a hypervirulent ST23 strain.

Using *in vitro* killing assays and Bliss analysis, we show that a 1:1 Loop and Spear cocktail did not improve bacterial suppression compared to Loop alone and instead exhibits multiplicity-of-infection–dependent antagonism, consistent with competition for a shared CPS receptor. Sequential dosing revealed strong order dependence: treatment with Spear followed by Loop qualitatively altered growth trajectories in a subset of cultures, whereas the reverse order or repeated dosing, provided little additional benefit.

Resistance profiling showed that mutants resistant to both phages predominantly carried mutations in capsule synthesis or export genes, whereas Spear specific resistance was associated with mutations in *fkpA*, encoding a periplasmic chaperone for outer membrane protein biogenesis. Adsorption assays confirmed that capsule associated mutations abolished Loop attachment, while Δ*fkpA* mutants retained Loop binding, supporting CPS to be a primary receptor with Spear additionally requiring an FkpA-dependent secondary receptor.

In a *Galleria mellonella* infection model, a capsule-mutant resistant isolate showed reduced virulence, and only sequential therapy with Spear followed by Loop improved survival beyond monotherapy. These findings show that receptor sharing can render phage cocktails antagonistic and highlight sequential, order-aware regimens as a strategy to exploit resistance–virulence trade-offs while limiting the emergence of double-resistant mutants.

## Introduction

*Klebsiella pneumoniae* is a major cause of healthcare-associated and community-acquired infections, including pneumonia, bloodstream and wound infections, and is a prominent driver of multidrug-resistant (MDR) Gram-negative sepsis [1–4]. Hypervirulent K1 ST23 lineages are of particular concern because they combine high levels of capsule-mediated immune evasion and tissue invasion with increasing antimicrobial resistance, leading to severe invasive disease [5–7]. Escalating carbapenem and extended-spectrum β-lactam resistance in *K. pneumonia* has created an urgent need for therapeutic alternatives that can either replace or potentiate antibiotics [8,9].

Bacteriophages are being actively explored as such alternatives, with multiple reports of successful *Klebsiella* phage therapy *in vitro*, in animal models and in compassionate-use cases [10–14]. Phage cocktails, a combination of two or more phages, are usually favoured over single phages because they are expected to broaden host range, reduce the probability of resistance and, in some settings, provide synergistic killing when combined with antibiotics [15–17]. In practice, however, many cocktails are assembled empirically based on host-range screening, morphology and genomic diversity; receptor usage, infection kinetics and pharmacodynamic interactions between constituent phages are only rarely characterised in detail. As a result, although cocktails are widely assumed to be beneficial, relatively few studies systematically assess whether phage–phage interactions are synergistic, additive or antagonistic across relevant multiplicities of infection and dosing regimens [18,19].

A key factor likely to influence such interactions is receptor usage. *K. pneumoniae* phages frequently target capsular polysaccharides (CPS) as primary receptors, sometimes in combination with secondary receptors such as lipopolysaccharide or outer membrane proteins [20], and capsule loss or modification is a common route to phage resistance that can reduce virulence [21,22]. In previous work, we characterised three genomically unrelated phages, Shorty, Spear and Loop, that infect the same hypervirulent K1 ST23 strain and showed that all three rely on CPS as their primary receptor [23]. Structural and genetic analyses revealed shared K1-specific lyase domains in the Loop and Shorty tail spikes despite otherwise distinct genomic backbones, and adsorption assays demonstrated that capsule-deficient mutants generated under Loop selection became resistant to all three phages. These findings highlighted CPS as a common vulnerability but also raised concerns that phages selected solely for diversity of morphology or genome could nonetheless converge on the same primary receptor, making cocktails potentially fragile to single-step resistance.

While receptor-guided cocktail design has been proposed conceptually, there is still limited experimental work that follows the full chain from receptor usage through resistance genetics to *in vivo* efficacy for defined phage combinations. In particular, several important questions remain unresolved. First, for phages that share a primary capsule receptor, it is unclear whether simultaneous application will yield improved suppression of *K. pneumoniae* or whether competition for the same receptor might instead lead to neutral or antagonistic interactions. Second, most phage therapy reports describe single or repeated dosing, but virtually none systematically examine whether the order of sequential administration of different phages affects outcomes, even though sequential use is clinically realistic. Third, although capsule loss is a well-established resistance mechanism, less is known about how different resistance trajectories, such as capsule synthesis versus outer membrane protein maturation defects, shape cross-resistance between phages, virulence *in vivo* and the performance of different dosing regimens.

Here, using the two K1-specific phages Loop and Spear as a defined test pair, we address these questions by integrating resazurin-based killing assays, Bliss analysis of phage–phage interactions, sequential dosing experiments, resistance genotyping, adsorption measurements and a *Galleria mellonella* infection model. We show that, despite their genomic distinctness, Loop and Spear do not form a synergistic cocktail and can display MOI-dependent antagonism when applied simultaneously. By contrast, sequential therapy is beneficial only when Loop is administered after Spear. We further show that mutants resistant to both phages predominantly carry capsule synthesis or export defects, whereas resistance specific to Spear maps to *fkpA*, which encodes a periplasmic chaperone involved in outer membrane protein biogenesis. Finally, a capsule-mutant resistant isolate displayed reduced virulence in *Galleria*, and the *in vivo* therapy experiments reproduce the same order dependence observed *in vitro*.

## Materials & Methods

### Bacterial strains and phage

The phage host strain is a hypervirulent *K. pneumoniae* of K1 ST23 serotype, obtained from the University Hospital in Leipzig, Germany. The negative control strain is *E. coli* BL21 DE3. For all experiments, the strains were cultured in LB medium from single colonies at 37 °C. Phages ‘Spear’ and ‘Loop’ were isolated from sewage water and characterised as described in an earlier study [23].

### Double-agar overlay

For all the experiments, 0.7% (w/v) LB agar was melted and mixed with bacterial host culture at OD_600_=1.0 at 5% (v/v). The soft agar was poured onto an LB plate to solidify. Phage samples were spotted onto a solidified lawn and plaques counted. The plates were always incubated at 37 °C.

### Resazurin-based growth curves

Bacterial culture was cultured up to OD_600_=1.3, and upon reaching kept on ice. The culture was diluted 1:100 in LB medium prior to seeding. A 96-well microtiter suspension culture plate was used for the assay. The total well volume was 200 µL, comprised 90 µL of phage sample at varying concentrations, 90 µL of diluted bacterial culture and 20 µL of 10x resazurin solution in PBS (440 µM). In the negative control group, phage samples were substituted with LB medium. Each plate had 4 sterility control groups: phage samples Spear and Loop, LB medium, and resazurin in LB. The plates were incubated at 37° C 218 RPM, taking 570 nm and 600 nm absorbance measurements every 100 sec over 16 h. For sequential addition, the incubation was interrupted at 6 h mark, and the wells were spiked with 10 µL of 10^8^ PFU/mL phage sample; the incubation proceeded for an additional 16 h.

### Synergy-antagonism analysis

Bliss independence analysis was conducted on wells with both phages Spear and Loop added simultaneously. For fractional inhibition analysis, the observed readout was converted as follows:

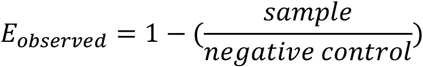

The Bliss predicted value was calculated as follows, where *S* and *L* are phages Spear and Loop, respectively:

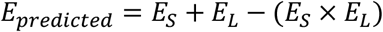

For the final Bliss excess values, the predicted value was subtracted from an observed value.

### Resistant variant generation

Bacterial culture at OD_600_=0.6 was infected with phages at a final MOI of 10 and overlayed via double-agar overlay. The plate was incubated at 37°C over a period of 16 h. Twenty-five distinct colonies were picked and streaked on a set of LB agar plates and incubated at 37°C for 16 h. Resulting colonies were propagated onto a new set of plates to produce uniform bacterial populations.

### Bacterial genome isolation

The genomes were isolated from using NEB Monarch^®^ Spin gDNA Extraction Kit according to manufacturer’s instructions. The whole-genome sequencing was performed by GENEWIZ via Illumina short-read sequencing (Microbe-EZ service).

### Variant Calling

Variant calling was done via *breseq* software v0.39.0 against raw FASTQ reads with chromosome-level assembled host genome used as a reference sequence [24]. The variants either reflecting a mixed population or producing an output with insufficient coverage were omitted from the final data set.

### Adsorption assay

Bacterial cultures were grown until an OD_600_=1.0 was reached. Phage samples at a titer of 10^9^ PFU/mL were added to the bacterial cultures at 10% (v/v). Bacteria-phage samples were incubated on ice for 20 min. The tubes were centrifuged at 16 000 × g, 4° C for 5 min. The supernatant was transferred, and a 10-fold dilution series was made. The titer of the supernatant was determined via double-agar overlay technique.

### Galleria mellonella survival assay

The *Galleria mellonella* survival assay was performed with final-instar larvae obtained from OwadoDajnia (Poland). Only larvae with a uniform cream colour and no visible melanisation were used. The larvae were divided into groups of 15 individuals, and each experiment was performed in three independent biological replicates. Larvae were infected with either the parental wild-type *K. pneumoniae* strain or its capsule-deficient mutant 310. Each larva received 3 µL of bacterial suspension corresponding to 10^5^ CFU/larva, injected into the hemocoel behind the last proleg. Phage treatment was administered 10 min after infection at MOI 100. For sequential treatment, the second phage dose was administered 8 h after the first dose. The experimental groups included bacteria-only controls, phage monotherapies with Spear or Loop, repeated phage treatments, and sequential combinations of both phages: Spear→Spear, Loop→Loop, Spear→Loop, and Loop→Spear. Non-injected larvae, PBS-injected larvae, and phage-only groups were included as controls. Larvae were incubated at 37°C for 96 h, and survival was recorded every 12 h. Larvae were considered dead when they showed melanisation and no response to physical stimulation.

Survival data were analysed in R using Kaplan–Meier survival curves. Differences between groups were assessed using log-rank tests with Benjamini–Hochberg correction for multiple comparisons. Median survival, survival at defined time points, restricted mean survival time, and Cox proportional hazards models were used to compare treatment effects.

## Results

### Loop and Spear do not form a synergistic cocktail

As phages are often used as cocktails, we first asked whether combining Loop and Spear improves suppression of *K. pneumoniae* growth beyond the activity of the individual phages. We monitored bacterial metabolic activity over time using a resazurin-based 96-well assay in which cultures were exposed either to increasing MOIs of each phage alone or to a 1:1 mixture at the corresponding total MOIs. Phage interactions were quantified using a Bliss-independence framework (Figure 1).

**Figure 1.**
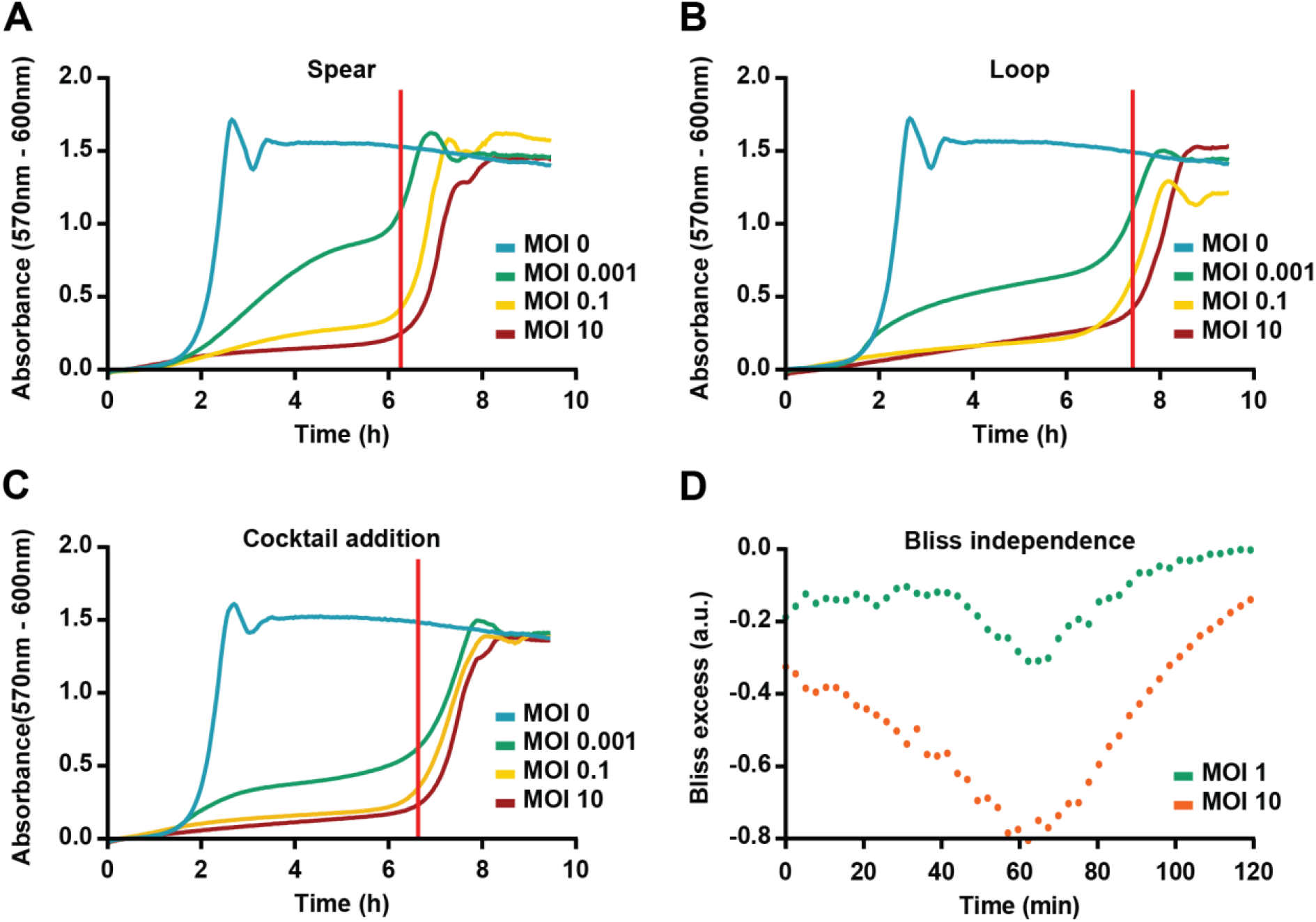
Loop–Spear cocktails suppress K. pneumoniae but show MOI-dependent antagonism. (A–C) Resazurin-based growth curves of the hypervirulent K1 ST23 K. pneumoniae strain exposed to different MOIs of phage Spear (A), phage Loop (B), or a 1:1 Spear–Loop cocktail (C). Metabolic activity (resazurin assay) was recorded over 10 hours. The vertical red line indicates the re-emergence of growth, most likely due to phage resistance. (D) Bliss-independence analysis of simultaneous Spear–Loop addition. Fractional inhibition values derived from the resazurin data were used to calculate Bliss-interdependence over 120 minutes at MOIs of 1 and 10. Bliss excess values around zero indicate additivity; negative values indicate antagonism. Data are representative of at least three independent experiments.

Spear alone showed clear MOI-dependent activity. At an MOI of 0.001, it delayed the increase in resazurin signal relative to the phage-free control, whereas MOIs of 0.1 and 10 suppressed bacterial growth for approximately 6-7 hours before regrowth emerged, probably due to phage resistance. Loop displayed a similar pattern but with slightly stronger suppression, with MOIs of 0.1 and 10 preventing bacterial growth for up to 7-8 hours, while an MOI of 0.001 again caused only delayed growth rather than complete inhibition. In the 1:1 cocktail condition, the killing curves essentially overlapped with those obtained for Loop alone at the corresponding total MOIs, indicating that the mixture did not improve suppression beyond that achieved by the more potent single phage and that the emergence of regrowth at later time points was not fully prevented by combining the two phages.

To assess whether the observed effects reflected synergy or antagonism, we performed Bliss-excess analysis over time. At a total MOI of 1, Bliss values remained close to or below zero, with a maximal deviation at approximately 60 minutes, consistent with mild antagonism. At a total MOI of 10, the Bliss excess became markedly negative at a similar time point, indicating stronger antagonistic interactions at higher phage concentrations. Because the magnitude of the effect increased with MOI, one possible explanation is that Loop and Spear compete for shared capsule binding sites, consistent with our previous finding that both phages primarily use the capsule as their receptor. Such competition could reduce the effective infection rate relative to that predicted from the activities of the individual phages. However, additional factors such as differences in secondary receptors, post-adsorption processes, or density-dependent changes in capsule accessibility, may also contribute to the observed antagonism and to its temporal variation.

Together, these results show that although both Loop and Spear effectively suppress *K. pneumoniae* growth at MOIs of 0.1 and above, combining them in a 1:1 cocktail does not enhance bacterial killing and instead results in concentration-dependent antagonism rather than synergy.

### Sequential application of Loop and Spear reveals order dependence

Because simultaneous addition of the Loop and Spear cocktail did not enhance killing and instead showed antagonism, we next examined whether sequential application of the two phages altered their combined effect on *K. pneumoniae* growth. Using the same resazurin-based 96-well assay, infections were initiated with a single phage, after which cultures were spiked at the indicated time point (after 6 hours, red line in Figure 2) with either the same phage or the alternate phage. Metabolic activity was then monitored over time.

**Figure 2.**
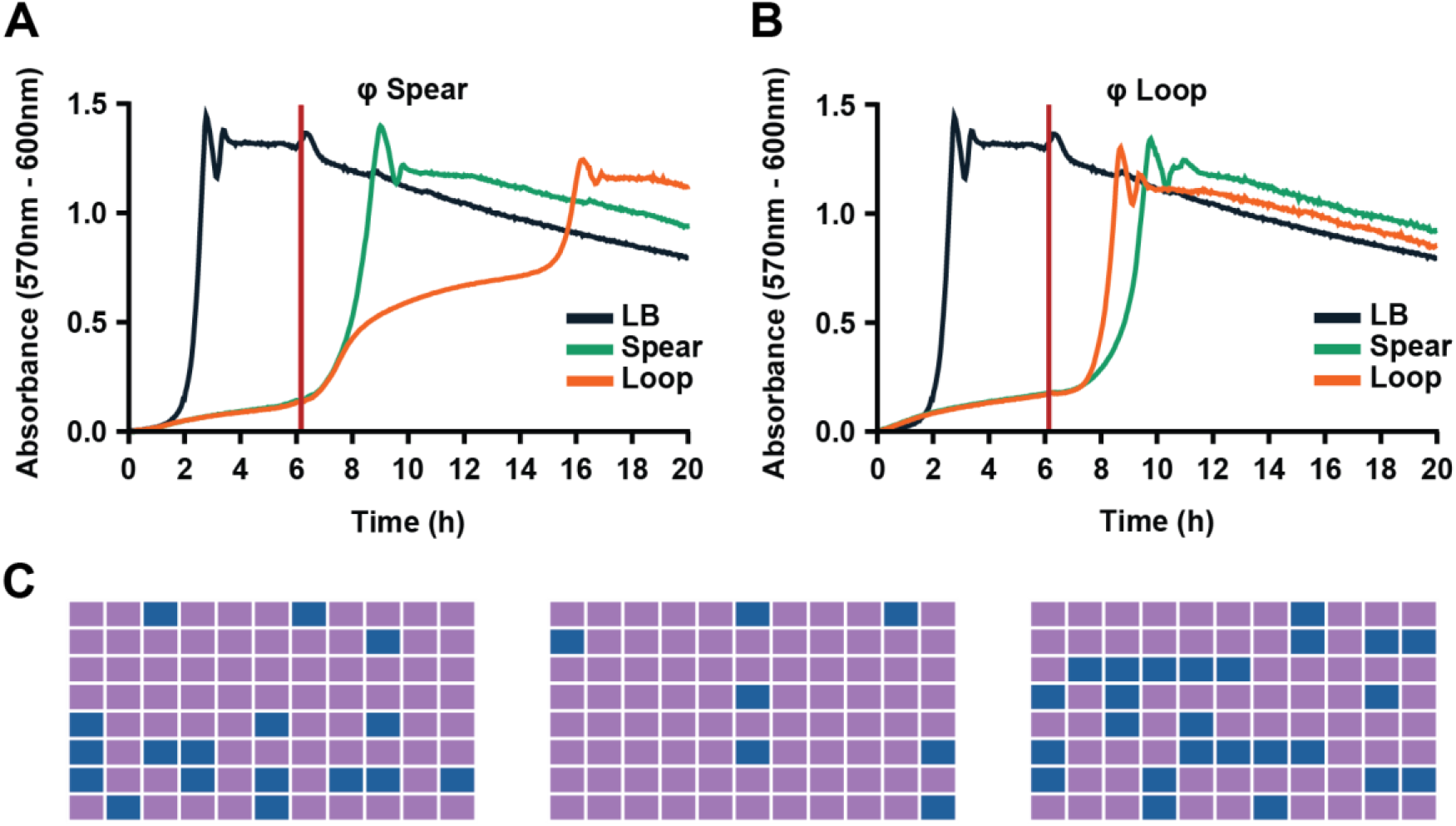
Sequential application of Loop and Spear reveals strong order dependence. (A, B) Resazurin-based growth curves for sequential dosing regimes. Cultures were infected in 96-well plates with phage Spear (A) or phage Loop (B) at MOI 1 at time 0, incubated for 6 hours, and then spiked with either the same phage or the alternate phage at 10^8 PFU/well. Metabolic activity was monitored for a further 14 hours. When Loop was given first (B), subsequent addition of Loop or Spear produced trajectories essentially indistinguishable from single-dose Loop. When Spear was given first (A), redosing with Spear had little effect relative to single-dose Spear, whereas adding Loop later produced a pronounced plateau in resazurin signal indicative of sustained growth suppression in a subset of cultures (yellow curve). (C) Heatmap of sequential-dosing outcomes across replicate wells. Each square represents a single well coloured according to endpoint phenotype (e.g. strong suppression vs regrowth). The Spear→Loop regime yielded a distinct “suppressed” pattern in on average 18.5% of wells (blue), whereas the other orders largely resembled the corresponding single-phage treatments.

When cultures were initially infected with Loop and subsequently received a second dose of Loop, the resazurin trajectories were largely unchanged relative to single-dose Loop treatment, indicating that repeated Loop administration did not further improve growth suppression under these conditions. Similarly, cultures initially treated with Loop and later spiked with Spear displayed growth dynamics closely resembling those of Loop alone, consistent with Loop dominating population suppression irrespective of subsequent Spear addition. Conversely, when cultures were first infected with Spear and later received a second dose of Spear, the growth dynamics differed little from single-dose Spear treatment, suggesting that repeated administration of the same phage has limited additional impact.

In contrast, when Spear was applied first and Loop was added later, we observed a pronounced plateauing in the resazurin signal following the second dose, consistent with substantially stronger inhibition of bacterial growth relative to Spear alone (Figure 2, yellow curve, panel A). A heatmap summary across experiments showed that this altered trajectory was not observed uniformly across well but occurred, on average, in 18.5% of cases. The remaining wells behaved similarly to Spear-only treatment, indicating that the effect is reproducible yet probabilistic rather than representing a deterministic shift in population behaviour.

These observations reveal a strong dependence of treatment outcome on phage order. Specifically, addition of Loop after Spear can qualitatively alter *K. pneumoniae* growth trajectories in a subset of cultures, whereas the reverse order and repeated dosing with the same phage produce little measurable change. These findings suggest that temporal order influences how emerging bacterial subpopulations interact with each phage.

### Phage selection reveals asymmetric cross resistance

Because the sequential dosing experiments indicated that prior exposure to Loop or Spear can influence subsequent treatment outcomes, we next examined how selection by each phage reshapes cross resistance patterns between them. *K. pneumoniae* cultures were exposed to either Spear or Loop, and individual colonies were isolated from regrowing cultures following phage treatment (“Spear resistance selection” or “Loop resistance selection” (Figure 3).

**Figure 3.**
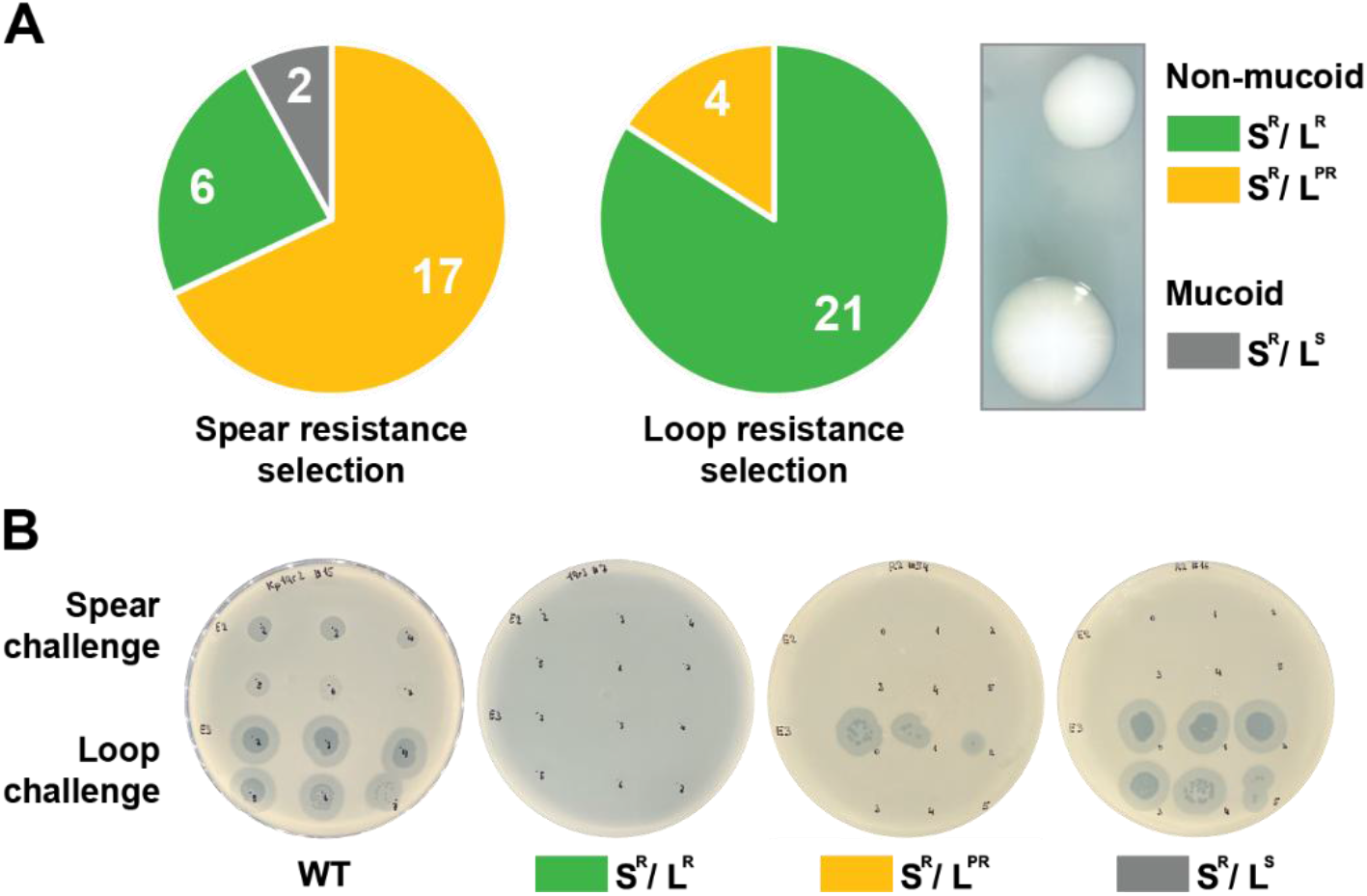
Phage selection generates asymmetric cross-resistance profiles. (A) Distribution of resistance phenotypes after selection with phage Spear or phage Loop. K. pneumoniae cultures were infected with phage Spear (left pie chart) or Loop (right pie chart) at MOI 10 in soft agar. After overnight incubation, 25 colonies were isolated from regrowing lawns, purified, and classified based on plaque assays with each phage as: Spear--and Loop-resistant (S^R^/L^R^, green), Spear-resistant and Loop-partially-resistant (S^R^/L^PR^, orange), or Spear-resistant and Loop-sensitive (S^R^/L^S^, grey). (B) Representative spot assays illustrating cross-resistance phenotypes. Wild-type (WT) and selected isolates from each class (S^R^/L^R^, S^R^/L^PR^, S^R^/L^S^) were spotted with serial dilutions of phage Spear (top row) or phage Loop (bottom row) on LB agar lawns. S^R^/L^R^ isolates lacked plaques for both phages, S^R^/L^PR^ isolates showed residual plaque formation for Loop only, and S^R^/L^S^ isolates remained fully susceptible to Loop while resistant to Spear. (C) Colony morphology of WT versus capsule-defective mutants. WT colonies display a hypermucoid appearance, whereas S^R^/L^R^ and S^R^/L^PR^ isolates show loss of mucoidy, consistent with capsule alteration or loss. S^R^/L^S^ mutants retain WT-like mucoid morphology.

Each isolate was subsequently tested for susceptibility to both phages and classified as Spear-resistant and Loop-resistant (S^R^/L^R^), Spear-resistant and Loop-partially-resistant (S^R^/L^PR^), or Spear-resistant and Loop-sensitive (S^R^/L^S^). Partial resistance was defined as the presence of a clear Loop mediated lysis zone on solid media despite the absence of a detectable effect on planktonic growth in liquid culture, which was caused by a drastic reduction capability of the phage to bind to the “partially resistant isolate” (L^PR^, Figure 4).

**Figure 4.**
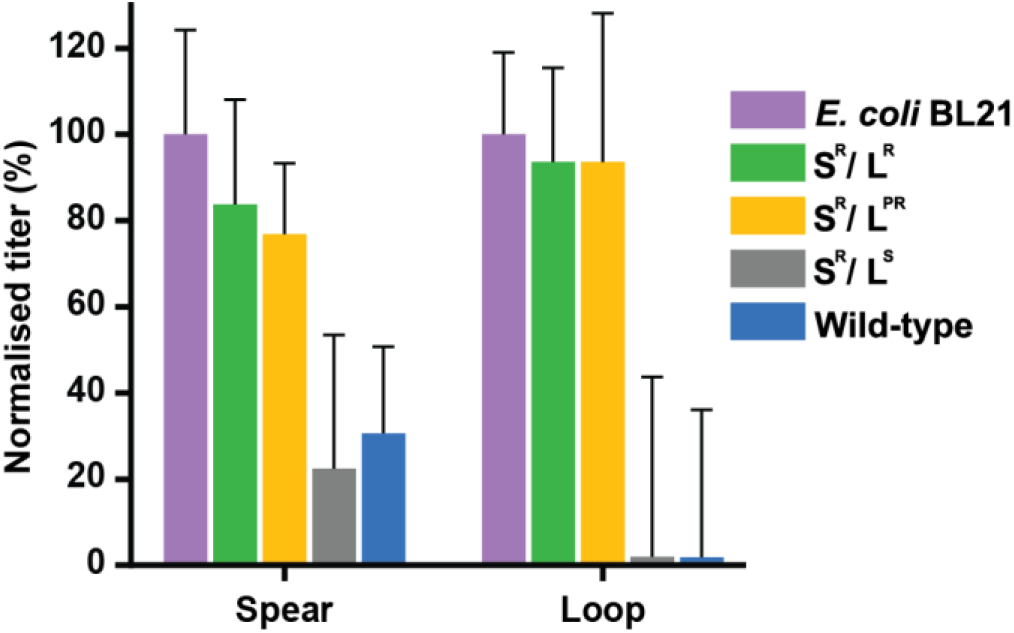
Capsule, but not fkpA, mutations abolish phage binding. Normalized Spear and Loop adsorption to WT and phage-resistant K. pneumoniae isolates. Determination of unbound phages to *E. coli* negative control, S^R^/L^R^, S*R*/L^PR^, S^R^/L^S^ and WT bacteria. Bars show mean normalized titers ± SD from three independent experiments.

Following Spear selection, most of the 25 recovered isolates were either fully (L^R^, n = 6) or partially resistant to Loop (L^PR^, n = 17), whereas 2 isolates remained fully sensitive to Loop (“L^S^”). In contrast, Loop selection yielded only S^R^/L^R^ (21 of 25) and S^R^/L^PR^ (4 of 25) isolates, and no Spear-sensitive mutants were recovered under these assay conditions. This asymmetry suggests that Loop preferentially selects mutants with reduced susceptibility to both phages, whereas Spear selection still allows the emergence of isolates that remain susceptible or only partially resistant to Loop. This observation is consistent with the stronger capsule dependence and higher adsorption efficiency previously described for Loop and aligns with the sequential dosing experiments, where cultures initially exposed to Spear remained susceptible to subsequent Loop treatment in approximately 18.5% of cases (see previous sub-section).

Phenotypically, S^R^/L^PR^ and S^R^/L^R^ isolates derived from both selection regimes consistently lost the hypermucoid colony morphology, whereas S^R^/L^S^ isolates retained the mucoid appearance of the wild type strain. Given our previous finding that the capsule serves as the primary receptor for Loop and contributes substantially to Spear adsorption, the association between loss of mucoidy and reduced phage susceptibility suggested capsule alteration or loss may represent a major resistance mechanism. To identify the underlying genetic changes directly, we therefore performed Whole Genome Sequencing (WGS) of the resistant isolates to establish the precise mutations involved and their role in resistance.

### Capsule associated mutations drive cross resistance, whereas *fkpA* mutations confer Spear specific resistance

Because Spear and Loop selection generated distinct resistance profiles and capsule associated phenotypes, we next asked which genetic changes underlie double resistance and Spear specific resistance. We performed whole-genome sequencing (WGS) and variant calling on 22 Loop generated variants (classified as S^R^/L^R^ or S^R^/L^PR^), as well as on the two S^R^/L^S^ isolates (Table 1).

**Table 1.**
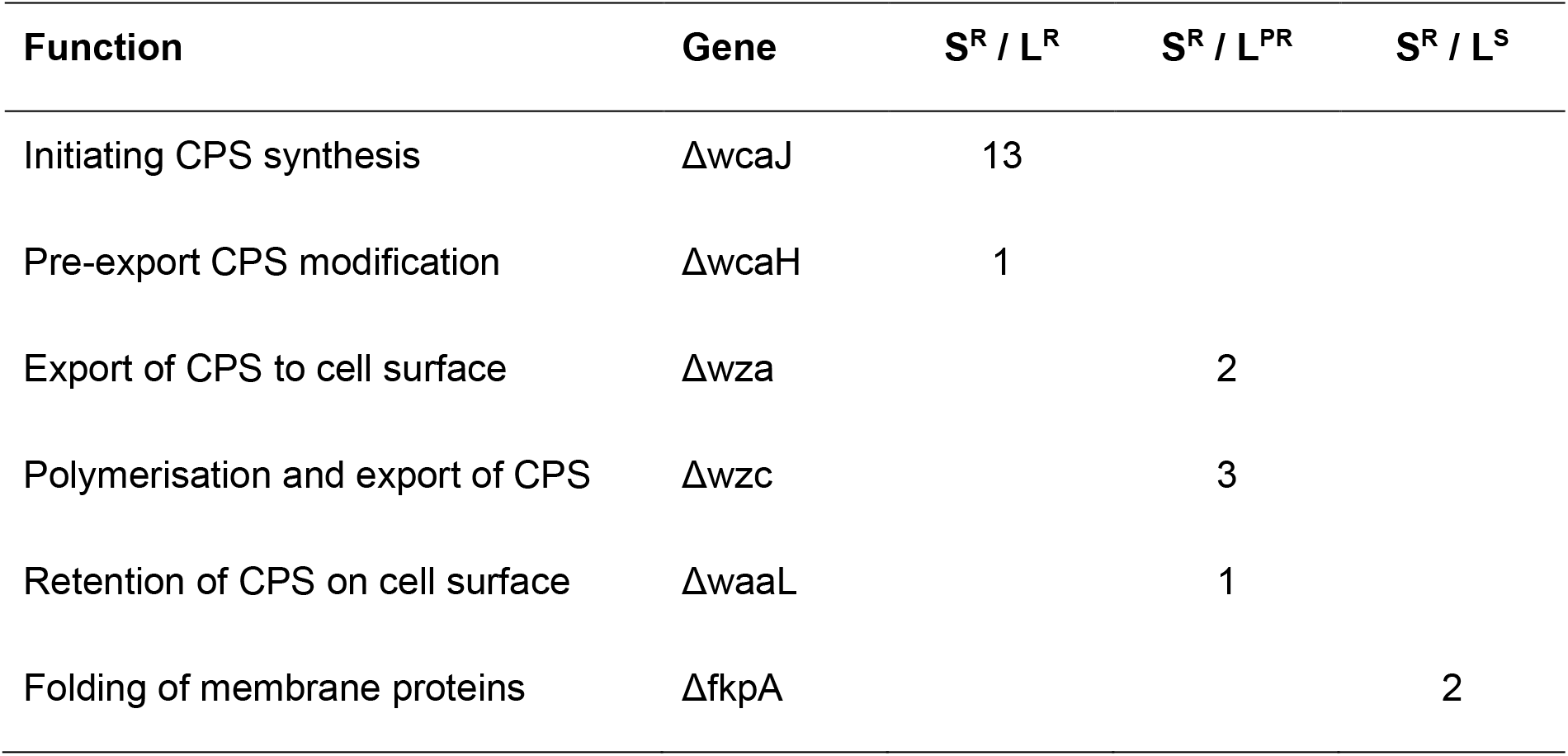
Capsule biosynthesis/export and fkpA mutations cause distinct resistance phenotypes. Summary of genes mutated in K. pneumoniae isolates resistant to phage Spear or Loop, grouped by resistance class: SR/LR (Spear--and Loop-resistant), SR/LPR (Spear-resistant and Loop-partially-resistant), and SR/LS (Spear-resistant and Loop-sensitive). Genes are organised by function along the capsule polysaccharide (CPS) pathway and outer membrane protein biogenesis. Early CPS synthesis genes (wcaJ and wcaH) are mutated almost exclusively in SR/LR isolates, indicating that disruption of the initial steps of capsule biosynthesis is the dominant route to double resistance. CPS export and polymerisation genes (wza, wzc) and CPS retention gene waaL are mutated in SR/LPR isolates, consistent with altered rather than abolished capsule display and partial Loop resistance. SR/LS isolates carry mutations only in fkpA, encoding a periplasmic chaperone for outer membrane protein folding and insertion, with no detectable capsule-pathway mutations.

| Function | Gene | S<sup>R</sup> / L<sup>R</sup> | S<sup>R</sup> / L<sup>PR</sup> | S<sup>R</sup> / L<sup>S</sup> |
| --- | --- | --- | --- | --- |
| Initiating CPS synthesis | $\Delta wcaJ$ | 13 | | |
| Pre-export CPS modification | $\Delta wcaH$ | 1 | | |
| Export of CPS to cell surface | $\Delta wza$ | | 2 | |
| Polymerisation and export of CPS | $\Delta wzc$ | | 3 | |
| Retention of CPS on cell surface | $\Delta waaL$ | | 1 | |
| Folding of membrane proteins | $\Delta fkpA$ | | | 2 |

Among mutants resistant to both phages (S^R^/L^R^), 13 of 14 carried mutations in *wcaJ*, which initiates CPS synthesis, whereas one isolate carried a mutation in *wcaH*, a gene implicated in capsule modification prior to export. These findings indicate that disruption of the earliest steps of capsule biosynthesis is the dominant route to double resistance in this dataset. By contrast, S^R^/L^PR^ isolates, which were resistant to Spear and only partially resistant to Loop, carried mutations in genes required for CPS export and surface polymerisation: two in *wza* (export of CPS to the surface), three in *wzc* (capsule polymerisation and export), and one in *waaL*, which is involved in CPS retention at the cell surface. These genotypes are consistent with altered rather than completely abolished capsule display in the partially resistant isolates. Together with the observed loss of mucoidy observed in S^R^/L^R^ and S^R^/L^PR^ isolates, these findings support the conclusion that both phages share CPS as a primary receptor and that different points of disruption along the CPS pathway gives rise to the spectrum ranging from full to partial resistance. However, functional analyses will be required to confirm the precise effects of each mutation on capsule structure and abundance.

In contrast, the two Spear resistant but Loop sensitive mutants (S^R^/L^S^) did not carry detectable mutations in capsule synthesis or export genes. Instead, both carried mutations in *fkpA*, which encodes a periplasmic chaperone involved in folding and insertion of outer membrane proteins (OMPs). Given that these *fkpA* mutants remain susceptible to Loop, whose infection is primarily CPS-dependent, while being resistant to Spear, our data suggests that Spear relies on an fkpA-dependent outer membrane protein as a secondary receptor downstream of CPS binding. We observed structural alignment of Spear tail fiber (CDS_0006) with the phage λ tip attachment protein J indicating that a potential receptor protein could be the OMP LamB (Supplementary Figure 1) [25].

### Capsule but not *fkpA* mutations impair Loop binding

To determine how the identified mutations affect the initial step of infection, we quantified Spear and Loop attachment to wild-type *K. pneumoniae* and representative S^R^/L^R^, S^R^/L^PR^, and S^R^/L^S^ mutants using an adsorption assay (Figure 4).

Cultures were incubated separately with Spear and Loop, after which bacterial cells were pelleted, and the remaining phage titer in the supernatant was quantified and normalized to the negative control (*E. coli*), in which no specific attachment is expected.

As anticipated, both phages showed efficient attachment to the wild-type host, with a low normalized titer in the supernatant compared to the *E. coli* control, indicating that most phage particles had been removed from the supernatant through binding to the true bacterial host. In contrast, both S^R^/L^R^ and S^R^/L^PR^ mutants displayed normalized titers close to the negative control, indicating a strong reduction or near-complete loss of Loop attachment. These findings align with the capsule synthesis and export defects identified in S^R^/L^R^ and S^R^/L^PR^ and indicate that disruption of the capsule pathway directly impairs binding, thereby explaining the observed resistance phenotypes in liquid culture.

By contrast, the S^R^/L^S^ mutants, which are resistant to Spear but remain susceptible to Loop, showed adsorption levels comparable to those of the wild type strain, with efficient removal of phage from the supernatant. Because these isolates carry *fkpA* mutations rather than mutations affecting CPS synthesis or export, this result supports the model that Loop relies exclusively on CPS for host recognition. Whereas Spear, although primarily binding to CPS, additionally depends on an fkpA-dependent outer membrane protein, likely LamB, as a secondary receptor. Disruption of *fkpA* abolishes Spear infection but not attachment, while leaving both Loop binding and infection largely unaffected. Meanwhile, capsule associated mutations eliminate attachment and infection by both phages.

Overall, the adsorption assays establish a direct functional link between the identified resistance mutations, receptor usage, and phage susceptibility phenotypes. Capsule synthesis and export mutations in S^R^/L^R^ and S^R^/L^PR^ mutants largely abolish Loop attachment, whereas *fkpA*-mediated Spear resistance leaves Loop binding intact.

### Sequential phage therapy remains order dependent in *Galleria mellonella*

Having established that sequential exposure to Spear and Loop produces order-dependent effects in vitro, we next tested whether this asymmetry translated into an *in vivo* infection setting. To this end, we used a *Galleria mellonella* survival model and compared infections caused by the parental WT *K. pneumoniae* strain and the capsule-deficient S^R^/L^R^ mutant 310. We then evaluated the therapeutic activity of Spear and Loop given either as monotherapies or as sequential regimens in which the second phage was administered 8 h after the first dose (Figure 5).

**Figure 5.**
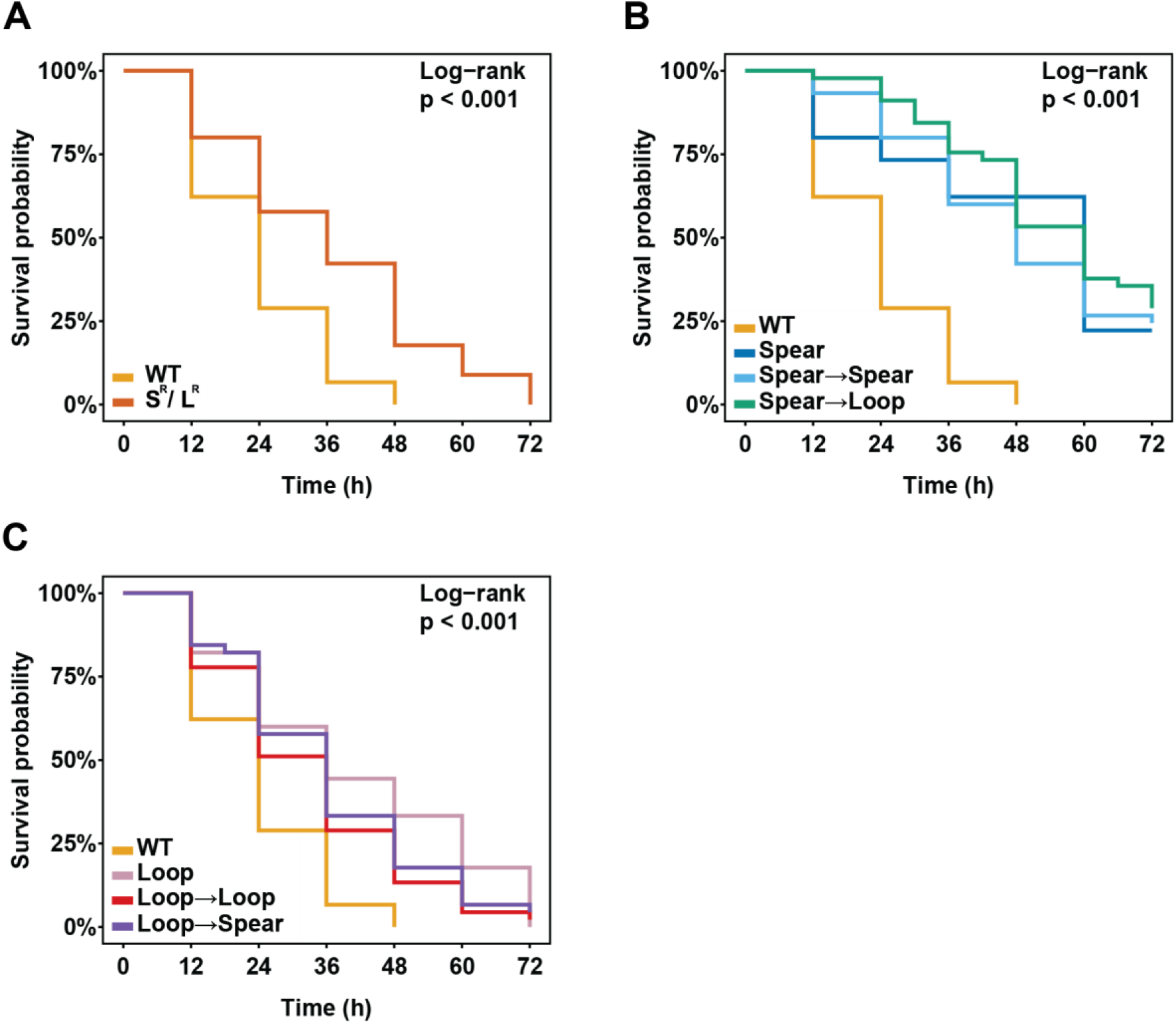
Sequential Spear→Loop therapy improves larval survival in vivo and remains dependent on treatment order. (A) Kaplan–Meier survival curves comparing larvae infected with the parental WT K. pneumoniae strain and the capsule-deficient S^R^/L^R^ mutant 310. (B) Survival of larvae infected with WT and treated with regimens initiated with Spear: Spear monotherapy, repeated Spear dosing, or sequential Spear→Loop treatment. (C) Survival of larvae infected with WT and treated with regimens initiated with Loop: Loop monotherapy, repeated Loop dosing, or sequential Loop→Spear treatment. Survival was monitored for 72 h and analysed using Kaplan–Meier curves and log-rank tests.

Infection with WT caused rapid larval mortality, with a median survival of 24 h and no larvae surviving to 72 h. In contrast, infection with 310 delayed disease progression, increasing median survival to 36 h and extending restricted mean survival time from 23.7 h to 36.8 h. Although 310 remained lethal under the conditions tested, the delayed mortality is consistent with reduced virulence relative to the parental strain and supports the role of the capsule as a major determinant of *K. pneumoniae* pathogenicity.

Phage treatment improved survival in a regimen dependent manner. Spear monotherapy provided the strongest single-phage effect, increasing median survival to 60 h and resulting in 22.2% survival at 72 h. Loop monotherapy was less protective, with a median survival of 36 h and no larvae surviving to the endpoint. Repeated Spear dosing did not substantially improve outcome relative to Spear alone, whereas the Spear→Loop sequence produced the most favourable survival profile among the tested regimens, with 28.9% survival at 72 h and the highest restricted mean survival time of 54.0 h. However, this improvement over Spear monotherapy did not remain statistically significant after correction for multiple comparisons.

The reciprocal sequence produced a different outcome. Loop→Spear failed to improve survival relative to Loop monotherapy and resulted in only 4.4% survival at 72 h. Direct comparison of the two reciprocal sequential regimens confirmed a significant order-dependent effect, with Spear→Loop outperforming Loop→Spear. These findings mirror the asymmetry observed in vitro: prior exposure to Spear creates a treatment context in which subsequent Loop administration provides additional benefit, whereas the reverse order does not confer comparable protection.

## Discussion

In this study, we used two previously characterized K1-specific *K. pneumoniae* phages, Loop and Spear, to dissect how shared receptor usage, resistance evolution and dosing regimen shape the performance of phage therapy candidates from *in vitro* assays to an *in vivo Galleria mellonella* infection model. Our main findings are that (i) simultaneous Loop–Spear cocktails are not synergistic and can be antagonistic despite phage diversity; (ii) sequential dosing is directionally effective, with benefits only when Loop is added after Spear; (iii) resistance to both phages is largely mediated by capsule synthesis and export defects, whereas Spear-only resistance maps to *fkpA* mutations; and (iv) at least one capsule-mutant resistant isolate is less virulent in *Galleria*, while the *in vivo* therapy experiments recapitulate the same order dependence observed *in vitro*. Together, these results argue that phage receptor usage and dosing order are critical parameters for designing robust therapies and that empirical cocktails based solely on phage diversity may be suboptimal or even counterproductive.

Simultaneous application of Loop and Spear did not enhance bacterial killing beyond the more potent single phage and even showed concentration (MOI)-dependent antagonism in Bliss analyses, despite the phages’ genomic distinctness and differences in receptor binding protein sequence and structure. This observation contrasts with the common assumption that combining multiple phages will almost automatically broaden host range and suppress resistance, an expectation often supported by case reports or small series rather than systematic pharmacodynamic evaluation. In our system, the most logical explanation for antagonism is competition for a shared primary receptor, the K1 capsule, at high local phage densities, which reduces effective adsorption relative to Bliss expectations. However, we cannot rule out additional contributors such as different secondary receptors, interference at post-adsorption steps or density-dependent changes in capsule accessibility, and these possibilities warrant dedicated follow-up experiments. The key point is that morphological and genomic diversity alone do not guarantee an effective cocktail; instead, receptor overlap and infection kinetics determined whether the combination was neutral, beneficial or antagonistic.

Sequential dosing experiments revealed that the temporal order of phage administration can qualitatively change infection outcomes. *In vitro*, repeating the same phage or adding Spear after Loop had little additional effect, whereas starting with Spear and adding Loop later converted a subset of otherwise surviving cultures into strongly suppressed ones. This directional effect was probabilistic rather than deterministic but reproducible, occurring in a consistent minority of wells. Importantly, the Galleria experiments recapitulated this order dependence: Spear→Loop treatment improved larval survival beyond either monotherapy or simultaneous dosing, whereas Loop→Spear did not provide a clear benefit over Loop alone. To our knowledge, such a matched *in vitro*/*in vivo* demonstration of order dependence for phage–phage combinations has not been reported, as most therapeutic reports emphasize dose, route and duration rather than the sequence in which individual phages are given. Our data suggest that, at least for phages sharing a primary receptor, sequential regimens need to be considered explicitly alongside cocktail versus single-phage choices.

By sequencing and phenotyping resistant isolates, we connected these phenotypic patterns to specific genetic routes of resistance. Most mutants resistant to both phages (S^R^/L^R^) carried mutations in *wcaJ* or *wcaH*, which initiate or modify capsule synthesis, whereas partially Loop-resistant mutants (S^R^/L^PR^) had mutations in *wza, wzc* or *waaL*, affecting capsule export and retention. These findings are consistent with our previous work identifying capsule as the primary receptor for both Loop and Spear and with broader literature showing that capsule-deficient or capsule-altered mutants frequently arise under phage selection and can exhibit attenuated virulence. Adsorption assays confirmed that these capsule-pathway mutations largely abolished Loop binding, mechanistically explaining double and partial resistance. In contrast, Spear-only resistant mutants that remained Loop-sensitive (S^R^/L^S^) lacked detectable CPS-pathway mutations but carried mutations in *fkpA*, encoding a periplasmic chaperone involved in folding and translocation of outer membrane proteins. Loop adsorption to these *fkpA* mutants remained comparable to wild type, indicating that Loop’s CPS-dependent infection is preserved whereas Spear-specific step is compromised. A parsimonious model is that Spear, but not Loop, additionally requires an *fkpA*-dependent outer membrane protein as a secondary receptor after capsule engagement, likely corresponding to LamB from distant structural homology search. Although its identity and its direct role in Spear infection remain to be demonstrated experimentally.

The *in vivo* experiments support and extend this mechanistic picture. Infection of *G. mellonella* with the wild-type K1 strain resulted in rapid larval killing, whereas infection with a representative capsule mutant resistant isolate led to reduced killing and prolonged survival, indicating that at least some phage-resistant pathways impose virulence costs *in vivo*. This is consistent with studies in *Klebsiella* and other Gram-negative pathogens showing that capsule or envelope modifications that confer phage resistance can impair colonisation, immune evasion or tissue invasion. From a therapeutic perspective, this trade-off is favourable; even when phage therapy does not fully eradicate the pathogen, it may shift the population towards phenotypes that are both more susceptible to host defences and less likely to cause severe disease. Overlaying this with the sequential dosing results suggests that a rational regimen could first use Spear to perturb the population and then apply Loop at a time when capsule-dependent infection is still possible, thereby reducing the likelihood of generating double-resistant mutants while preferentially selecting less virulent capsule-deficient variants.

Our findings are broadly consistent with an emerging view that phage therapy should move from empirical mixing of “cocktails” to a more pharmacology-like framework that accounts for receptor usage, resistance pathways and dosing dynamics. At the same time, they challenge the simplistic notion that more phages in a cocktail are always better. In our system, simultaneous application of two capsule-targeting phages neither improved *in vitro* killing nor enhanced *in vivo* survival compared to the better single phage and risked driving bacteria towards double resistance, whereas a carefully chosen sequential regimen exploited their mechanistic complementarity. These results do not argue against cocktails per se, especially when phages are chosen to avoid receptor overlap, but they underscore that receptor-informed design and explicit testing of sequential orders should be standard in preclinical evaluation.

Several limitations need to be acknowledged. First, our resistance genotyping focused on Loop-generated variants and a small number of Spear-selected S^R^/L^S^ mutants; mixed populations and low coverage prevented a comprehensive view of Spear-selected clones, and additional work will be required to define the full spectrum of resistance routes under different regimens. Second, the proposed f*kpA*-dependent secondary receptor for Spear remains a hypothesis; targeted construction of clean *fkpA* knockouts and complementation, along with systematic outer membrane proteomics and adsorption assays, will be necessary to identify the target protein and confirm the two-step receptor model. Third, the *Galleria* model captures key aspects of virulence and treatment but cannot fully recapitulate mammalian immunity, pharmacokinetics or tissue distribution. Future work in mammalian models will be essential to test whether the same order dependence and resistance–virulence trade-offs hold in more complex settings.

Despite these limitations, our study advances the conceptual and practical basis for designing phage therapies. It shows that receptor sharing can turn diverse phages into antagonistic partners when administered together, that sequential administration can exploit directional effects to improve outcomes, and that mapping resistance genotypes onto receptor use and virulence can reveal which phage combinations and dosing orders are most likely to benefit patients rather than accelerate the emergence of problematic double-resistant mutants. More broadly, it argues for integrating receptor-guided phage selection, quantitative combination analysis (including Bliss-type metrics) and sequential dosing experiments into preclinical pipelines, so that clinical regimens can be built on mechanistic understanding rather than empirical mixtures alone.

## Author Contributions (according to CRediT: https://credit.niso.org/)

**ZS**: Conceptualization, Investigation, Formal analysis, Methodology, Validation, Visualization, Writing—original draft, Writing – review & editing. **PO**: Investigation, Data curation, Validation. **BG**: Data curation, Validation, Methodology, Writing—review & editing. **SL**: Conceptualization, Visualization, Methodology, Writing—original draft, Writing – review & editing. **BL**: Conceptualization, Formal analysis, Supervision, Resources, Visualization, Writing – review & editing.

## Data availability

All variant raw reads together with the annotated wild-type host genome are available in European Nucleotide Archive (ENA) under the following project accession: PRJEB114347.

