## Supplementary Figure 1 for "Sequential Use of Two Capsule-Targeting Klebsiella Phages Reveals Order-Dependent Efficacy and Distinct Resistance Pathways *in vitro* and *in vivo*"

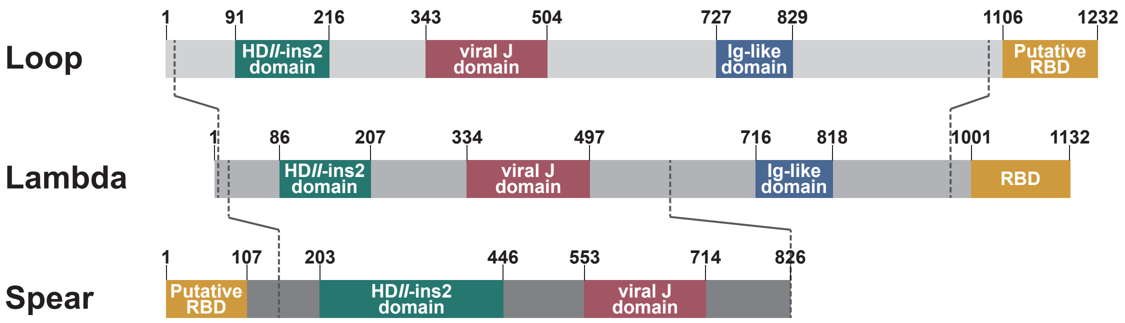


Supplementary figure 1. Structural homology of Spear tail fiber J (CDS_0005) and Loop tail fiber J (CDS_0009). Both proteins align to Lambda phage's tip attachemnt protein J (UniProtKB: P03749). HDII-ins2 and viral J domains are conserved across all three proteins, with an additional Ig-like domain conserved between fibers of Lambda and Loop.
